# Genome-wide screens reveal shared and strain-specific genes that facilitate enteric colonization by *Klebsiella pneumoniae*

**DOI:** 10.1101/2023.08.30.555643

**Authors:** Bettina H Cheung, Arghavan Alisoltani, Travis J Kochan, Marine Lebrun-Corbin, Sophia H Nozick, Christopher MR Axline, Kelly ER Bachta, Egon A Ozer, Alan R Hauser

## Abstract

Gastrointestinal (GI) colonization by *Klebsiella pneumoniae* is a risk factor for subsequent infection as well as transmission to other patients. Additionally, colonization is achieved by many strain types that exhibit high diversity in genetic content. Thus, we aimed to study strain-specific requirements for *K. pneumoniae* GI colonization by applying transposon insertion sequencing to three classical clinical strains: a carbapenem-resistant strain, an extended-spectrum beta-lactamase producing strain, and a non-epidemic antibiotic-susceptible strain. The transposon insertion libraries were screened in a murine model of GI colonization. At three days post-inoculation, 27 genes were required by all three strains for colonization. Isogenic deletion mutants for three genes/operons (*acrA*, *carAB*, *tatABCD*) confirmed colonization defects in each of the three strains. Additionally, deletion of *acrA* reduced bile tolerance *in vitro*, while complementation restored both bile tolerance *in vitro* and colonization ability *in vivo*. Transposon insertion sequencing suggested that some genes were more important for colonization of one strain than the others. For example, deletion of the sucrose porin-encoding gene *scrY* resulted in a colonization defect in the carbapenemase-producing strain but not in the extended-spectrum beta-lactamase producer or the antibiotic-susceptible strain. These findings demonstrate that classical *K. pneumoniae* strains use both shared and strain-specific strategies to colonize the mouse GI tract.

**IMPORTANCE:** *Klebsiella pneumoniae* is a common cause of difficult-to-treat infections due to its propensity to express resistance to many antibiotics. For example, carbapenem-resistant *K. pneumoniae* (CR-Kp) has been named an urgent threat by the United States Centers for Disease Control and Prevention. Gastrointestinal colonization of patients with *K. pneumoniae* has been linked to subsequent infection, making it a key process to control in prevention of multidrug-resistant infections. However, the bacterial factors which contribute to *K. pneumoniae* colonization are not well understood. Additionally, individual strains exhibit large amounts of genetic diversity, begging the question of whether some colonization factors are strain-dependent. This study identifies the enteric colonization factors of 3 classical strains using transposon mutant screens to define a core colonization program for *K. pneumoniae* as well as detecting strain-to-strain differences in colonization strategies.

## INTRODUCTION

As multidrug-resistant bacteria continue to pose a looming threat to our ability to treat infections, alternative methods for controlling disease burden—such as infection prevention—are increasingly important. For *Klebsiella pneumoniae*, a highly multidrug-resistant bacteria, gastrointestinal (GI) colonization is a risk factor for subsequent infection. Patients are prone to infections caused by the strains they carry and may also spread them to other hospitalized patients^1,2^. Thus, GI colonization by *K. pneumoniae* is an attractive target for infection prevention.

The genes required for *K. pneumoniae* gut colonization are not well understood. Studies of intestinal colonization are complicated by the fact that this species exhibits substantial genetic diversity. While the average *K. pneumoniae* genome contains about 5000-6000 genes, only ∼1700 are shared by most strains^3,4^. Studies have identified colonization factors for individual strains^5–7^, but whether *K. pneumoniae*’s genetic diversity gives rise to strain-to-strain differences in GI colonization strategies remains unclear.

Two broad categories of *K. pneumoniae* strains have been described: classical and hypervirulent. Of the classical strains, several have achieved global dominance and are referred to as high-risk clones. Two such high-risk clones are the ST45 and ST258 sequence types. ST45 strains frequently produce extended-spectrum beta-lactamases (ESBLs), making them resistant to most beta-lactams except carbapenems. Many ST258 strains produce carbapenemases, causing a significant portion of carbapenem-resistant infections worldwide^8^. While it is conceivable that the global success of these lineages has resulted in part from their capacity to better colonize the GI tract, how their colonization strategies might differ from those of antibiotic-susceptible strains has not been investigated.

Here, we selected three representative clinical classical strains of *K. pneumoniae* to investigate GI colonization by lineages of varying epidemicity and antibiotic resistance. We used transposon insertion sequencing to identify genes in each strain that were required for GI colonization in a mouse model. We found that a subset of genes was required by all three strains, but each strain relied on a much larger set of additional genes that were dispensable for the other strains. Thus, the set of genes, and therefore presumed GI colonization strategies, vary substantially across phylogenetically distinct *K. pneumoniae* strains.

## RESULTS

### Three clinical strains of *K. pneumoniae* with distinct genetic and phenotypic characteristics

We selected three clinical isolates of *K. pneumoniae* representative of strains with varying levels of epidemic spread and antibiotic resistance. First, we chose CRE-166, a carbapenem-resistant strain of the ST258 high-risk clone with the *bla*_KPC_ gene, which was isolated from bronchioalveolar lavage. Second, we selected Z4160, an ESBL producer isolated from the bloodstream with both a widespread ESBL gene (*bla*_CTX-M-15_) and an epidemic sequence type (ST45). Third, we chose KPN46, a non-epidemic, antibiotic-susceptible bloodstream isolate (ST433). The sizes of the CRE-166, Z4160, and KPN46 genomes were 6.00, 5.56, and 5.63 Mb, respectively. The core genome shared between them was 4.96 Mb (4,662 coding sequences [CDS]), leaving CRE-166, Z4160, and KPN46 with 1.04, 0.63, 0.69 Mb (1183, 623, and 749 CDS) of accessory genetic content (Figure 1). Thus, CRE-166, Z4160, and KPN46 represented three clinical strains with phenotypic and genomic diversity suitable for subsequent studies of *K. pneumoniae* GI colonization.

**Figure 1.**
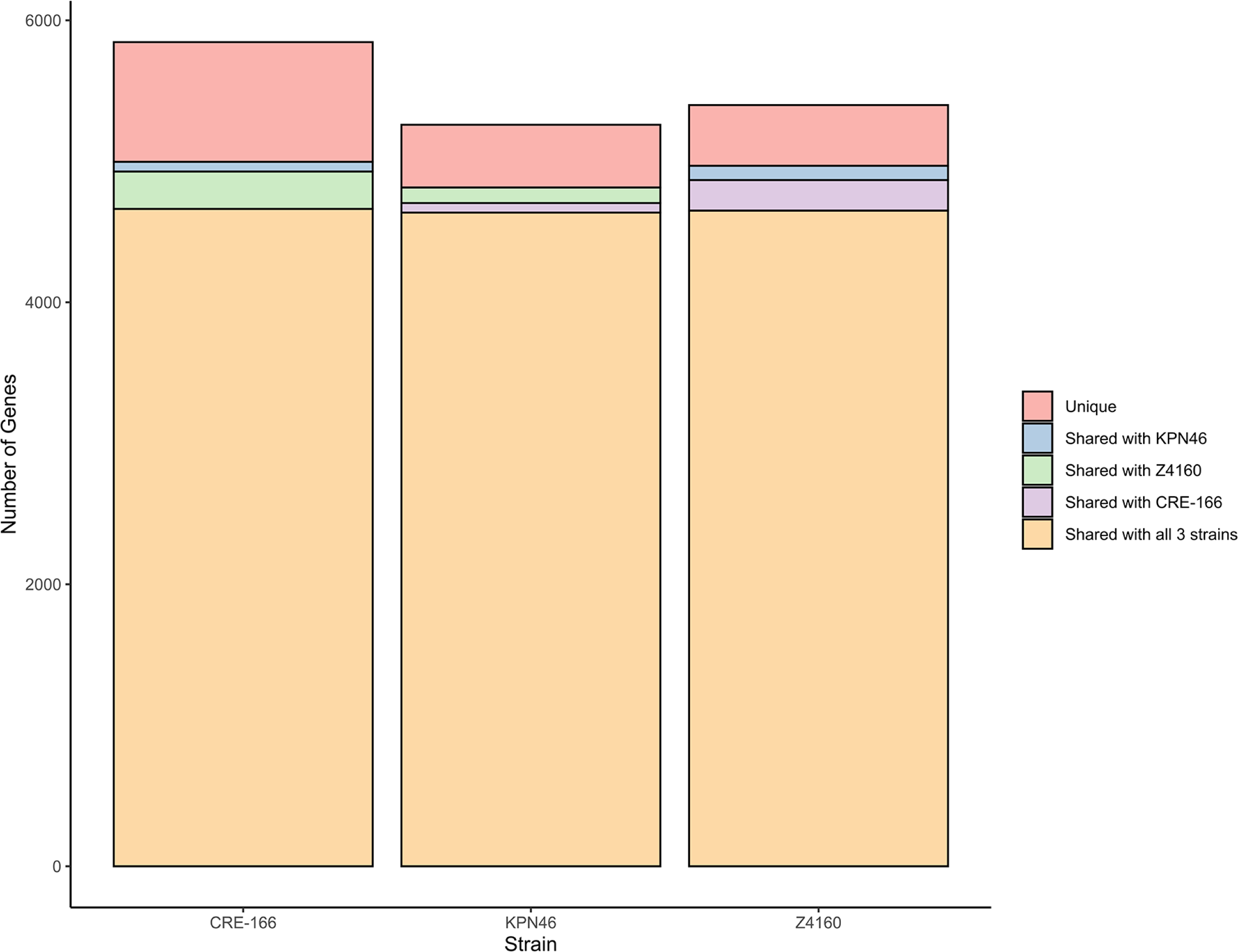
Unique and shared coding sequences between 3 strains of *K. pneumoniae*. The software program Spine was used to identify coding sequences with at least 85% homology in the strains indicated.

### A clinically relevant murine model of GI colonization

To model hospitalized patients receiving antibiotics and to induce robust fecal shedding of *K. pneumoniae* in C57BL/6 mice, we administered an antibiotic regimen prior to oral gavage with bacteria. Vancomycin, one of the most highly utilized antibiotics in the United States^9^, was administered through daily intraperitoneal injection for 5 days (350 mg/kg, equivalent to a human dose of 1 g/day)^10^ (Figure 2A). In contrast to other screens for GI colonization factors in which antibiotics were administered through drinking water and throughout the screen^5–7^, we opted for injections to control dosage to each mouse and to allow for precise adjustments in future studies. We also ceased administration prior to *K. pneumoniae* inoculation to investigate colonization after antibiotic exposure. Following the last dose of vancomycin, 10^8^ colony forming units (CFU) of CRE-166 were inoculated by orogastric gavage. *K. pneumoniae* was then selectively cultured from feces by plating on lysogeny broth (LB) agar supplemented with carbenicillin, an antibiotic to which all *K. pneumoniae* are resistant^4^. Culture of feces on this medium prior to inoculation with *K. pneumoniae* yielded no colonies, confirming specificity for experimentally introduced *K. pneumoniae* (Supplemental Figure 1A). These conditions supported 10^10^ CFU/g fecal shedding of *K. pneumoniae* in the first week followed by shedding of 10^7^ CFU/g for at least 60 days post-gavage in both male and female mice (Figure 2B). At Day 14, no CFU could be detected in the lung, liver, or spleen, indicating there was no dissemination from the gut (Supplemental Figure 1B). Additionally, throughout the infection, mice did not exhibit signs of illness, suggesting that they were colonized rather than infected.

**Figure 2.**
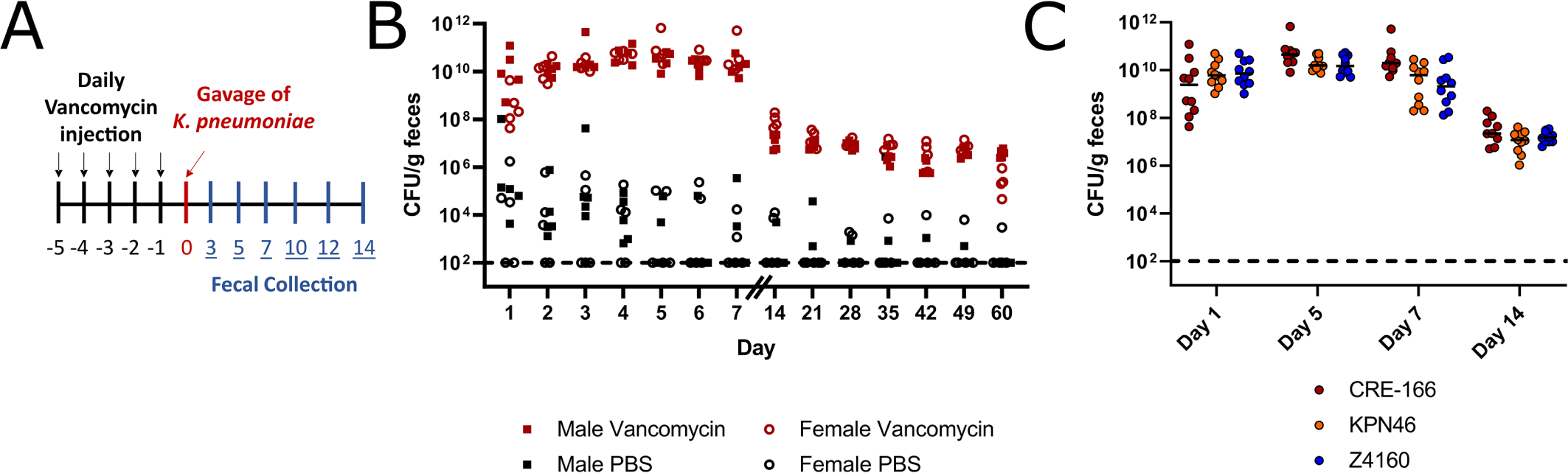
Mouse model of GI colonization with *K. pneumoniae*. (A) Schematic of the *in vivo* model. Mice were administered 5 days of 350 mg/kg intraperitoneal vancomycin injections before orogastric gavage with 10^8^ CFU *K. pneumoniae*. Fecal samples were collected after gavage and CFU enumerated. (B) Fecal burden of CRE-166 following gavage into male (square) or female (circle) mice with (red) or without (black) vancomycin treatment prior to gavage. n = 5 for each group. (C) Fecal burdens of strains CRE-166, KPN46, and Z4160 in mouse model of GI colonization. n = 10 for each group. Line indicates median. Limit of detection was 10^2^ CFU/g feces, denoted by a dotted line.

To ensure these conditions would produce similar levels of GI colonization with KPN46 and Z4160, we individually inoculated CRE-166, KPN46, and Z4160 into mice. Fecal burdens were similar across 14 days (Figure 2C). These results indicated that the vancomycin-treated mice were a robust model of *K. pneumoniae* GI colonization.

To determine whether transposon insertion sequencing experiments would be informative in our model, we next ensured that mutants would not randomly drop out of the fecal output due to bottlenecks rather than colonization defects. To this end, we constructed a marked CRE-166 strain by inserting an apramycin-resistance cassette into the chromosomal Tn7 site. This marked strain did not have a growth defect in LB when compared to the parental strain (Supplemental Figure 2A). To mimic the presence of a single transposon mutant within the pool of total mutants in a screen, we spiked this marked strain into an inoculum at a ratio of 1:100,000 with the parental strain. Next, we measured the ratio of the marked strain to total *K. pneumoniae* recovered from the feces of the mice. At Day 3 post-gavage, the marked strain was still detectable, suggesting the absence of a bottleneck significant enough to bias transposon sequencing results (Supplemental Figures 2B & 2C). However, at subsequent timepoints, we failed to recover the marked strain from some of the mice, indicating greater bottlenecks. We therefore chose Day 3 post-gavage as the timepoint for transposon sequencing experiments.

### Generation of highly saturated transposon mutant libraries

To perform genome-wide screens for GI colonization factors, we generated transposon mutant libraries in all three *K. pneumoniae* strains. The transposon vector pSAM*erm* was modified to express hygromycin resistance (pSAM*hygSDM*) to allow for selection of transposition in all three strains. Libraries with over 145,000 mutants were generated. An initial assessment of library quality was performed by randomly selecting 32 colonies from each library and identifying transposon insertion sites with arbitrary PCR. Unique insertion sites for at least 26 colonies for each strain were successfully identified, and no colonies had more than one insertion site, indicating the libraries were of high quality.

### Screening the mutant libraries for GI colonization factors *in vivo*

We gavaged each of the three transposon mutant libraries into mice pre-treated with vancomycin. A portion of the inoculum was saved as the “input pool.” At Day 3, we collected fecal pellets, or the “output pools.” Input pool sequencing demonstrated that over 82% of coding sequences had at least one insertion, and on average, each gene had 5 insertions. Coverage was distributed across chromosomes (Supplemental Figure 3), confirming that all libraries were well-saturated.

Insertion site sequencing reads were then processed using a modified version of the previously described ESSENTIALS pipeline^11^. We first analyzed the input pools to identify “essential genes” required for the bacteria to grow in LB. A total of 487 genes were identified as essential in all three strains, but a substantial number of genes were essential in only one or two strains (Figure 3A, Supplemental Table 1). Between 14-18% of CDS in each strain were found to be essential, comparable to the estimates of 11-17% reported for other strains of *K. pneumoniae*^5,12–14^.

**Figure 3.**
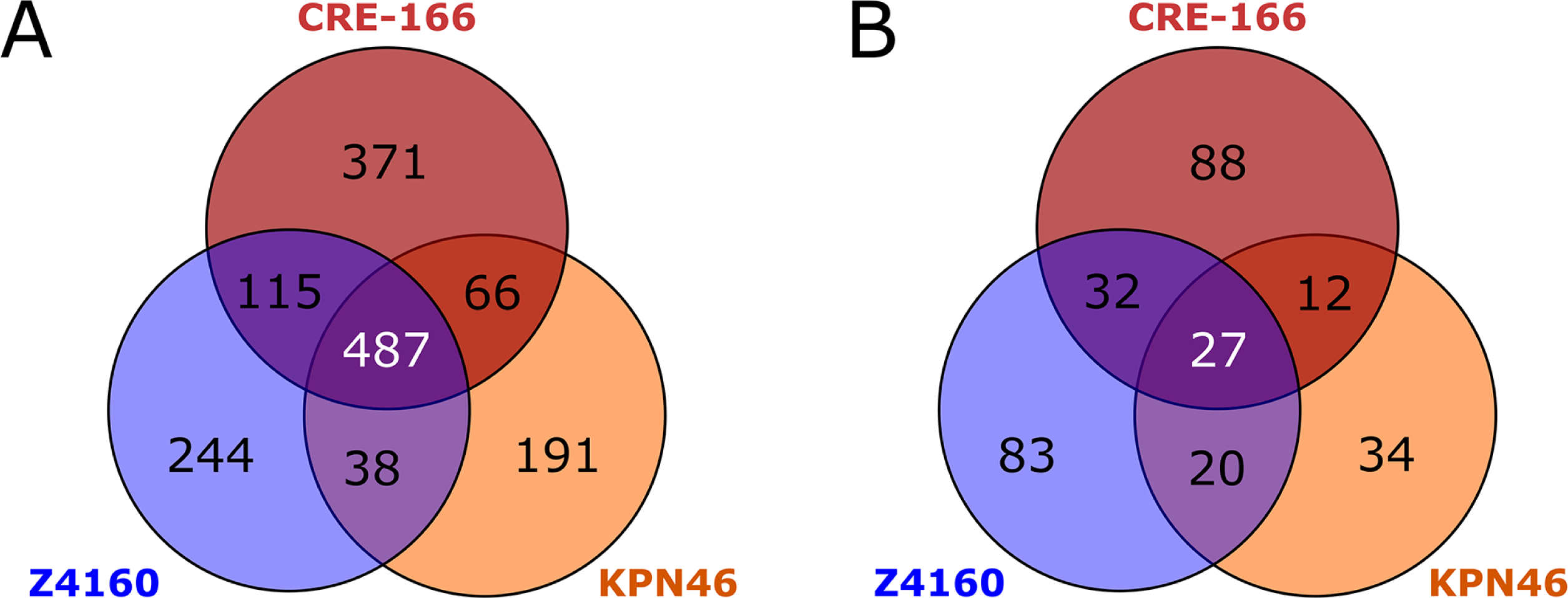
Genes required for growth in LB and for GI colonization. Transposon mutant libraries were screened in a mouse model of GI colonization. n = 3 for inputs and outputs for each strain. (A) Shared and unique essential genes for each strain. (B) Shared and unique genes for which transposon insertions resulted in reduced fitness in the gut. Genes had a log_2_ fold change < −2 and a false discovery rate < 0.05.

To determine which genes each strain utilized for establishing GI colonization, we compared the total number of insertion reads per gene in the Day 3 output pool to those of the input pools. We focused on genes that had a less than −2 log_2_(fold-change) (logFC) in output vs. input insertion reads and a false discovery rate (FDR) less than 0.05 (Figure 4, Supplemental Table 2). Multidimensional scaling (MDS) plots of input and Day 3 output pools demonstrated that the input pools were closely related and distinct from the Day 3 output pools for each strain (Supplemental Figure 4), indicating our approach was technically robust.

**Figure 4.**
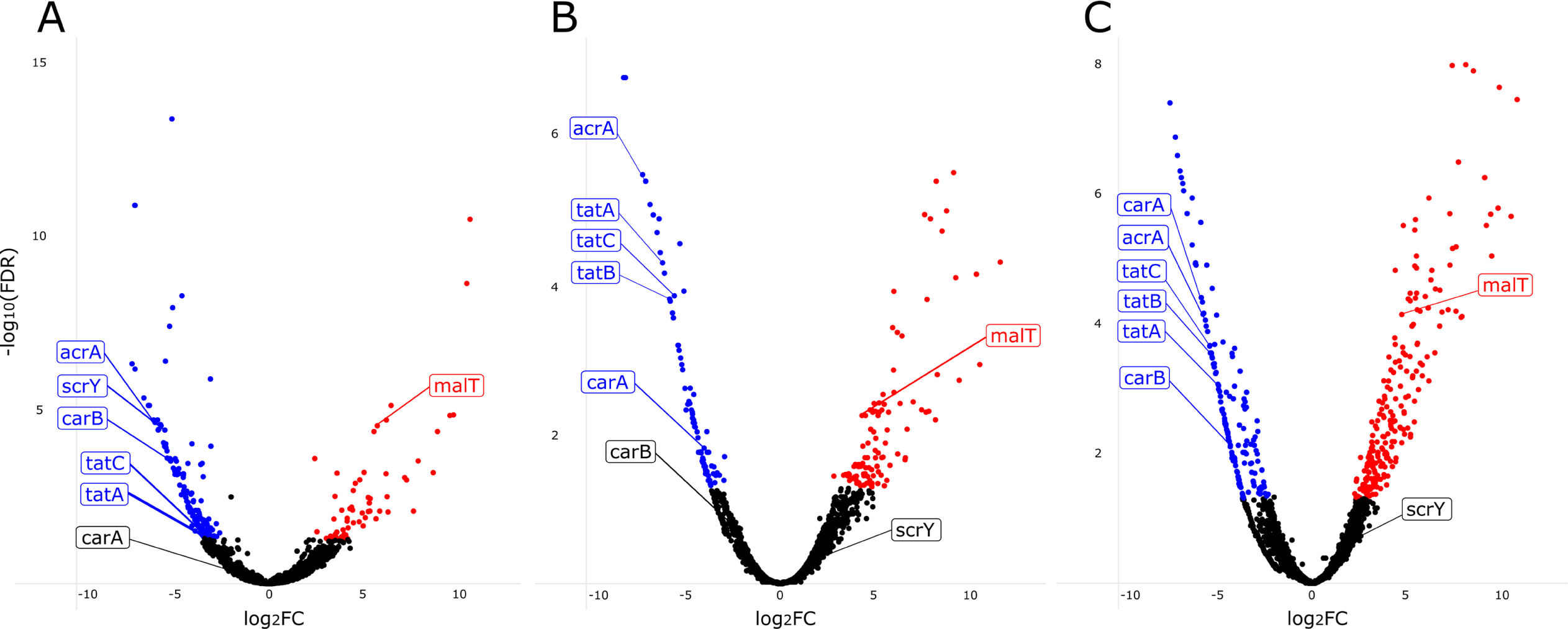
Volcano plots showing results of transposon insertion sequencing experiments in mice. Genes in which insertions were associated with decreased GI colonization (blue dots) and increased colonization (red dots) for (A) CRE-166, (B) KPN46, and (C) Z4160 are shown. Labeled points indicate targets that were chosen for the creation of isogenic mutants. FDR, false discovery rate; FC, fold change.

Twenty-seven genes were used by all three strains for GI colonization (Figure 3B). However, many genes were used by only one strain to establish colonization: 88 for CRE-166, 83 for Z4160, and 34 for KPN46. Intriguingly, most genes identified as important for colonization in at least one strain were present in all 3 strains. That is, only 22.6% of colonization genes for CRE-166, 3.3% for KPN46, and 6.8% for Z4160 were absent from the genomes of the other strains, suggesting that these strains mostly rely on shared genes to establish GI colonization, but use different sets of these genes for this purpose.

To determine whether these colonization genes were also found in broader populations of *K. pneumoniae* strains, we calculated a core genome (genes shared by 95% of strains) from a set of 323 previously described strains^4^. Upon comparison with this broader core genome, somewhat larger percentages of the colonization genes for each strain were now considered accessory genes: 25.8% for CRE-166, 16.1% for KPN46, and 15.4% for Z4160. However, most genes required for colonization by each strain were still genes shared across *K. pneumoniae* strains.

In addition to genes required for colonization, we also identified genes which, upon disruption with a transposon, conferred a colonization advantage (red points in Figure 4). There were 7 genes found to confer an advantage when disrupted in all 3 strains (Supplemental Table 2). Four were involved in maltose transport (*malT*, *malEFG*) while the other three had regulatory roles (*proQ*, *prc*, and *rspR*).

### Classification of GI colonization factors

To better understand the core colonization program in *K. pneumoniae*, we focused on the 27 genes important for colonization across all three strains (Table 1). As expected, we identified genes involved in anaerobic metabolism (e.g., *adhE, fnr, focA).* We also found genes involved in other metabolic pathways, including *mtlD* (mannitol-1-phosphate dehydrogenase) and *carAB*, which encode the subunits of carbamoyl phosphate synthase that are responsible for the first committed step in synthesis of pyrimidine and arginine^15^. *carA* was identified in two strains and *carB* in the remaining strain. *tatA* and *tatC* (folded protein secretion apparatus^16^) and *acrA* (efflux pump^17^) were also identified.

**Table 1.** Genes contributing to GI colonization in all 3 strains of *K. pneumoniae*.

| Gene | Annotation | log<sub>2</sub>(Fold change) |  |  |
| --- | --- | --- | --- | --- |
|  |  | CRE-166 | KPN46 | Z4160 |
| *aceE* | Pyruvate dehydrogenase E1 component | -4.412 | -5.504 | -5.440 |
| *acrA* | Multidrug efflux pump subunit | -5.906 | -6.935 | -5.512 |
| *adhE* | Aldehyde-alcohol dehydrogenase | -3.353 | -4.848 | -7.339 |
| *arcB* | Aerobic respiration control sensor protein | -5.342 | -3.821 | -6.018 |
| *bglY* | Beta-galactosidase | -6.948 | -2.896 | -5.749 |
| *cvpA* | Colicin V production protein | -4.611 | -4.449 | -6.031 |
| *cydA* | Cytochrome bd-I ubiquinol oxidase subunit 1 | -4.387 | -6.933 | -4.731 |
| *fnr* | Fumarate and nitrate reduction regulatory protein | -4.248 | -6.542 | -5.723 |
| *focA* | Formate Transporter | -6.486 | -7.092 | -6.679 |
| *glnA* | Glutamine synthetase | -3.667 | -3.755 | -4.313 |
| *miaA* | tRNA dimethylallyltransferase | -5.773 | -3.856 | -3.992 |
| *mtlD* | Mannitol-1-phosphate 5-dehydrogenase | -5.711 | -6.709 | -5.990 |
| *ompC* | Outer membrane porin C | -5.738 | -3.129 | -3.472 |
| *pal* | Peptidoglycan-associated lipoprotein | -4.172 | -6.183 | -4.662 |
| *pflA* | Pyruvate formate-lyase 1-activating enzyme | -5.617 | -7.979 | -6.821 |
| *pflB* | Formate acetyltransferase 1 | -4.766 | -5.270 | -4.077 |
| *pgi* | Glucose-6-phosphate isomerase | -5.404 | -5.665 | -4.507 |
| *ptsI* | Phosphoenolpyruvate-protein phosphotransferase | -3.088 | -6.240 | -7.061 |
| *purC* | Phosphoribosylaminoimidazole-succinocarboxamide synthase | -3.470 | -4.634 | -4.273 |
| *purH* | Bifunctional purine biosynthesis protein | -3.710 | -5.209 | -8.324 |
| *pykF* | Pyruvate kinase I | -4.431 | -4.701 | -3.025 |
| *setA* | Sugar efflux transporter A | -6.943 | -5.168 | -5.053 |
| *tatA* | Sec-independent protein translocase protein | -3.384 | -6.061 | -4.829 |
| *tatC* | Sec-independent protein translocase protein | -3.452 | -5.452 | -5.276 |
| *tolA* | Tol-Pal system protein | -5.977 | -5.965 | -4.284 |
| *yeiE* | HTH-type transcriptional activator | -5.445 | -4.572 | -4.522 |
| - | Hypothetical polysaccharide deacetylase | -5.327 | -4.906 | -3.985 |
| *carA* | Carbamoyl phosphate synthase small subunit | - | -3.917 | -5.662 |
| *carB* | Carbamoyl phosphate synthase large subunit | -5.267 | - | -4.220 |

\*For each strain, genes with log<sub>2</sub>fold change < -2 and FDR < 0.05 were compared. For the *carAB* two-gene operon, *carB* met these criteria for CRE-166 whereas *carA* met these criteria for KPN46 and Z4160.

To examine the pathways that strains relied on for colonization, we assigned Kyoto Encyclopedia of Genes and Genomes (KEGG) identifiers to all genes in each strain and determined which pathways were enriched among colonization hits. These pathways fell into a few broad categories: metabolism, antimicrobial resistance, protein secretion, and environmental sensing (Figure 5). Metabolic capacities played an important role in the ability of bacteria to colonize the gut, but individual pathways identified differed by strain. Additionally, the colonization factors for CRE-166 and KPN46 were enriched for two-component systems, which may have played a role in metabolic adjustments caused by environmental sensing in the GI tract. In terms of antimicrobial resistance pathways, two strains (CRE-166 and KPN46) were reliant on genes that conferred resistance to cationic antimicrobial peptides (CAMPs), which are released by colonic epithelium and are similar to microcins released by the microbiota. Thus, defense against host and microbiome factors is likely key to colonization by *K. pneumoniae*. KPN46 colonization factors were enriched for genes for resistance to beta-lactams, including efflux pumps, which play a role in the efflux of toxic compounds. Finally, protein export (the Tat secretion system) was enriched for KPN46 and Z4160, suggesting that secreted proteins may enhance colonization.

**Figure 5.**
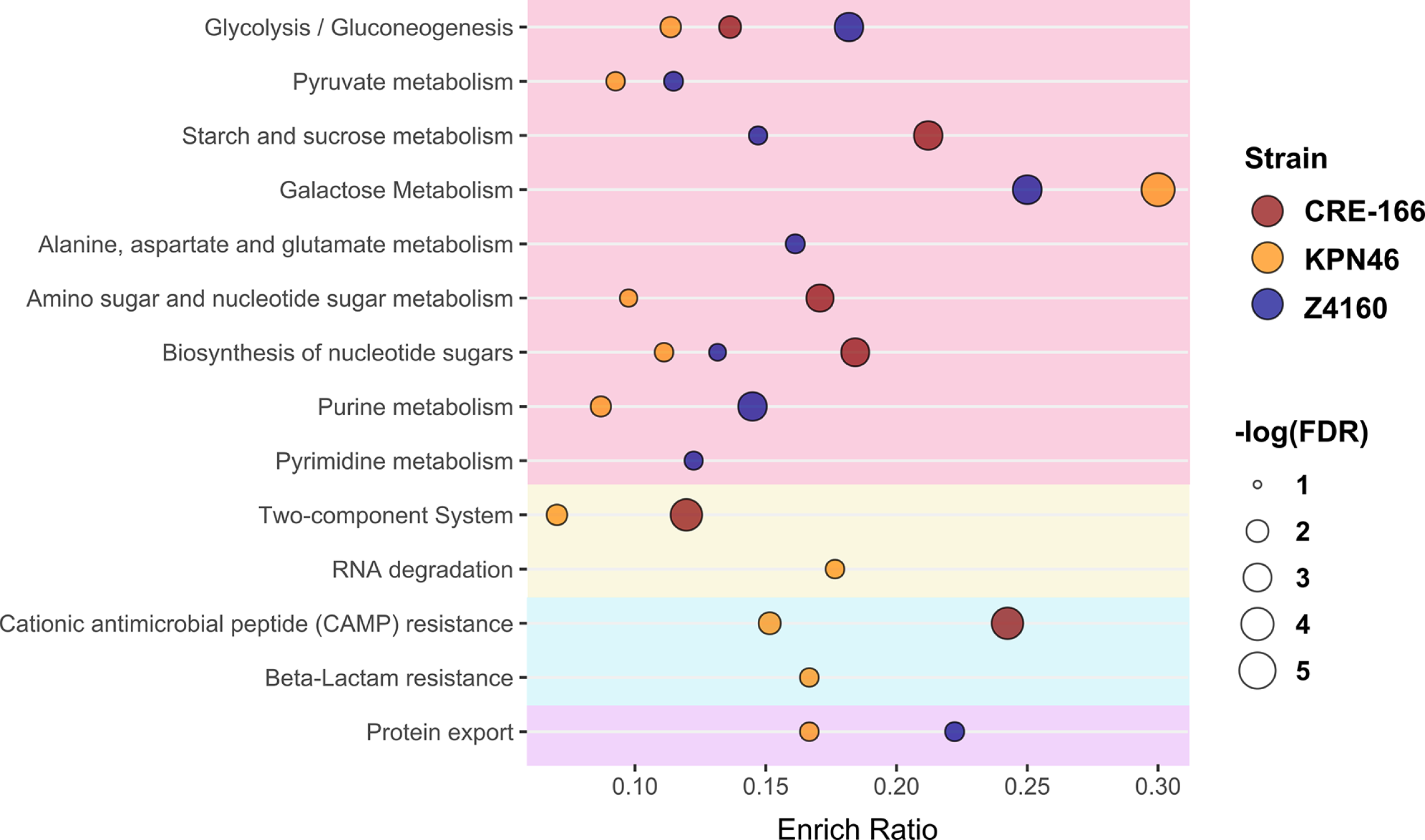
Pathways used for GI colonization in 3 strains of *K. pneumoniae*. Kyoto Encyclopedia of Genes and Genomes (KEGG) identifiers were assigned to all genes in the genomes of CRE-166, KPN46, and Z4160. The enrichment ratio (Enrich Ratio) was calculated as the ratio of genes in the target list belonging to the specified KEGG pathway to the total number of genes in the pathway in the genome. A hypergeometric test was used to determine false discovery rate (FDR), and FDR < 0.05 was considered significant.

### Validation of colonization genes using isogenic mutants

To validate our screen, we created isogenic mutants of 3 genetic loci—*acrA*, *carAB*, and *tatABCD*—required for GI colonization in all 3 strains (Table 1). These loci were chosen because they represent different functional groups: antimicrobial resistance, metabolism, and secretion. We generated isogenic mutants in which the coding sequence of the target was replaced by an apramycin-resistance cassette. We verified that these apramycin-resistant mutants did not have growth defects in LB when grown individually (Supplemental Figure 5). Of these mutants, only the *tatABCD* mutant had a slight defect in LB when competed against their marked (hygromycin-resistant) parental strains (Supplemental Figure 6). Then, we inoculated 1:1 mixtures of the marked parental strains and mutants into the mouse model of GI colonization and enumerated CFU in the feces at Day 3 (the screen timepoint) to calculate competitive indices (CI). To characterize the effects of these mutants at later timepoints, we also followed the fecal burdens to Day 14.

In all three strain backgrounds, *acrA* mutants displayed significant colonization defects at Day 3 (validating our screen) as well as beyond to Day 14 (Figure 6A-C). We constructed a complemented strain with an unmarked deletion of the *acrA* locus, inserting *acrA* along with its upstream region and a downstream apramycin-resistance cassette into the chromosomal Tn7 site. This complement rescued the colonization defect (Figure 6D).

**Figure 6.**
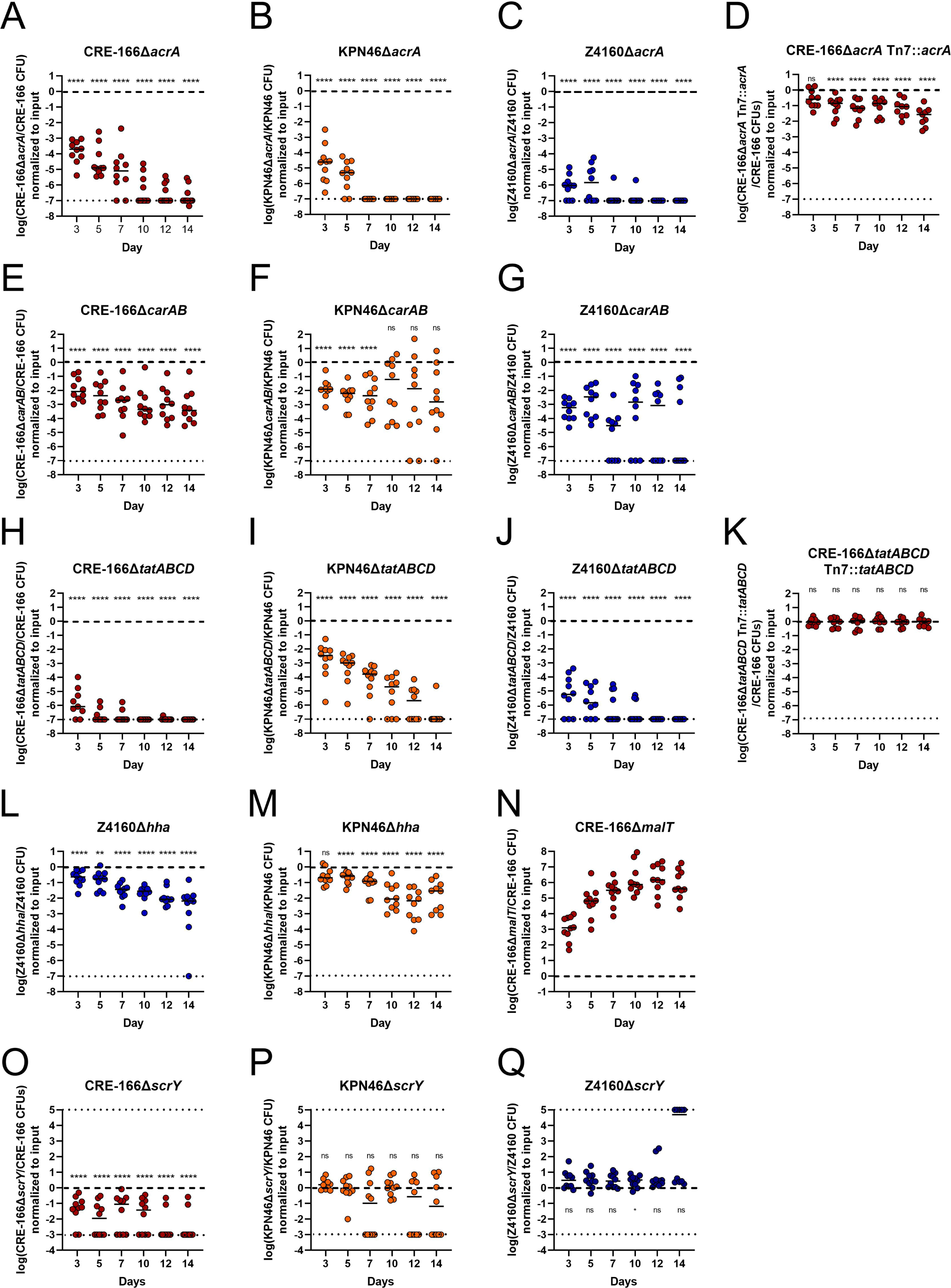
Competitive colonization between parent strains and isogenic mutants to validate genes identified in transposon insertion screens. Mice were treated with 5 days of vancomycin prior to gavage with 1:1 mixtures of marked parent strain (hygromycin-resistance cassette at the Tn7 site) and isogenic mutant (substitution of open reading frame with apramycin-resistance cassette [A-N] or an unmarked in-frame deletion [O-Q]) of the target gene(s). n = 10 for A-G and L-N. n ≥ 9 for H-K and O-Q. Asterisks denote significance by one-sample t-tests with Dunn correction where * p < 0.05, ** p < 0.01, *** p < 0.001, **** p < 0.0001, and “ns” indicates not significant. Limit of detection was a competitive index of 10^-7^ for A-N and 10^-3^ for O-Q, denoted with a dotted line. Log(competitive index) = 0, or equal recovered CFU of parental strain and mutant, is marked with a dashed line.

The *carAB* deletion mutants were similarly tested in competition with their parental strains. At Day 3, each *carAB* mutant exhibited colonization defect, continuing to Day 14 for CRE-166 and Z4160 (Figure 6E-G). For the KPN46 mutant, greater variability in CI was observed at later timepoints, suggesting the existence of a priority effect in the second week, during which mutants that initially established themselves tended to do very well while the others did progressively more poorly. Due to technical limitations, a *carAB* complement could not be constructed. However, we performed whole genome sequencing on all mutants, confirming that the CRE-166 and Z4160 mutants did not have off-site mutations likely to be responsible for observed phenotypes. For KPN46, sequencing indicated that two nonsynonymous mutations emerged during the course of mutant generation. We performed an *in vivo* competition experiment between two marked KPN46 parental strains—one with these off-site mutations and one without—to show that the mutations did not confer a colonization defect (Supplemental Figure 7). These data indicate that the *carAB* deletions are responsible for the colonization defects observed in Figure 6E-G.

Finally, deletion of the *tatABCD* operon also significantly decreased colonization capacities, both at Day 3 and throughout subsequent days (Figure 6H-J). Insertion of the *tatABCD* operon at the Tn7 site fully rescued the colonization defect (Figure 6K). Thus, we verified our ability to detect shared factors essential for GI colonization.

In addition to shared factors, we wanted to confirm strain-specific colonization factors. Hemolysin expression-modulating protein, encoded by *hha*, is a transcriptional regulator which scored as a hit in Z4160 but not in KPN46 and CRE-166. We generated *hha* deletion mutants in both Z4160 and KPN46, neither of which had *in vitro* growth defects (Supplemental Figure 5). At Day 3, the Δ*hha* mutant had a statistically significant colonization defect in Z4160 (Figure 6L). In KPN46, the Δ*hha* mutant had a slightly less severe colonization defect that was not statistically significant (Figure 6M).

As a second strain-specific factor, we selected *scrY*, which encodes for a sucrose porin, and which was identified in our screen as a colonization factor for CRE-166 but not KPN46 or Z4160. We created in-frame deletions to preserve the remainder of the operon downstream from *scrY* and found that CRE-166Δ*scrY* but not KPN46Δ*scrY* or Z4160Δ*scrY* had a colonization defect (Figure 6O-Q). Together, these data indicate that colonization factors may differ in their importance from strain to strain.

Finally, we also selected one target that exhibited a colonization *advantage* upon disruption. We chose *malT*, the transcriptional regulator for maltose uptake and metabolism. A *malT* deletion mutant in CRE-166 did not have a growth advantage in LB (Supplemental Figure 5), but this deletion conferred a substantial colonization advantage over the parent strain *in vivo* (Figure 6N), indicating our screen was also valid for detection of genes that confer colonization advantages.

### Deletion of *acrA* reduces resistance to ox bile

We further explored how one of our colonization genes, *acrA,* may contribute to GI colonization. As *acrA* encodes a component of an efflux pump that contributes to bile resistance in other GI pathogens^18^, we tested whether our *acrA* mutant was more sensitive to bile. At 2 and 24 hours after inoculation into 10% bile, the marked parental strain grew significantly better than the CRE-166Δ*acrA*::AprR mutant (Figure 7). We also generated a CRE-166Δ*acrA* mutant in which the apramycin-resistance cassette was removed from the Δ*acrA* allele; this mutant also showed a growth defect in bile. A complemented strain (CRE-166Δ*acrA* Tn7::*acrA*) generated from this mutant showed that complementation rescued resistance to bile. Together, these data suggest that *acrA* supports *K. pneumoniae* GI colonization by providing resistance to bile.

**Figure 7.**
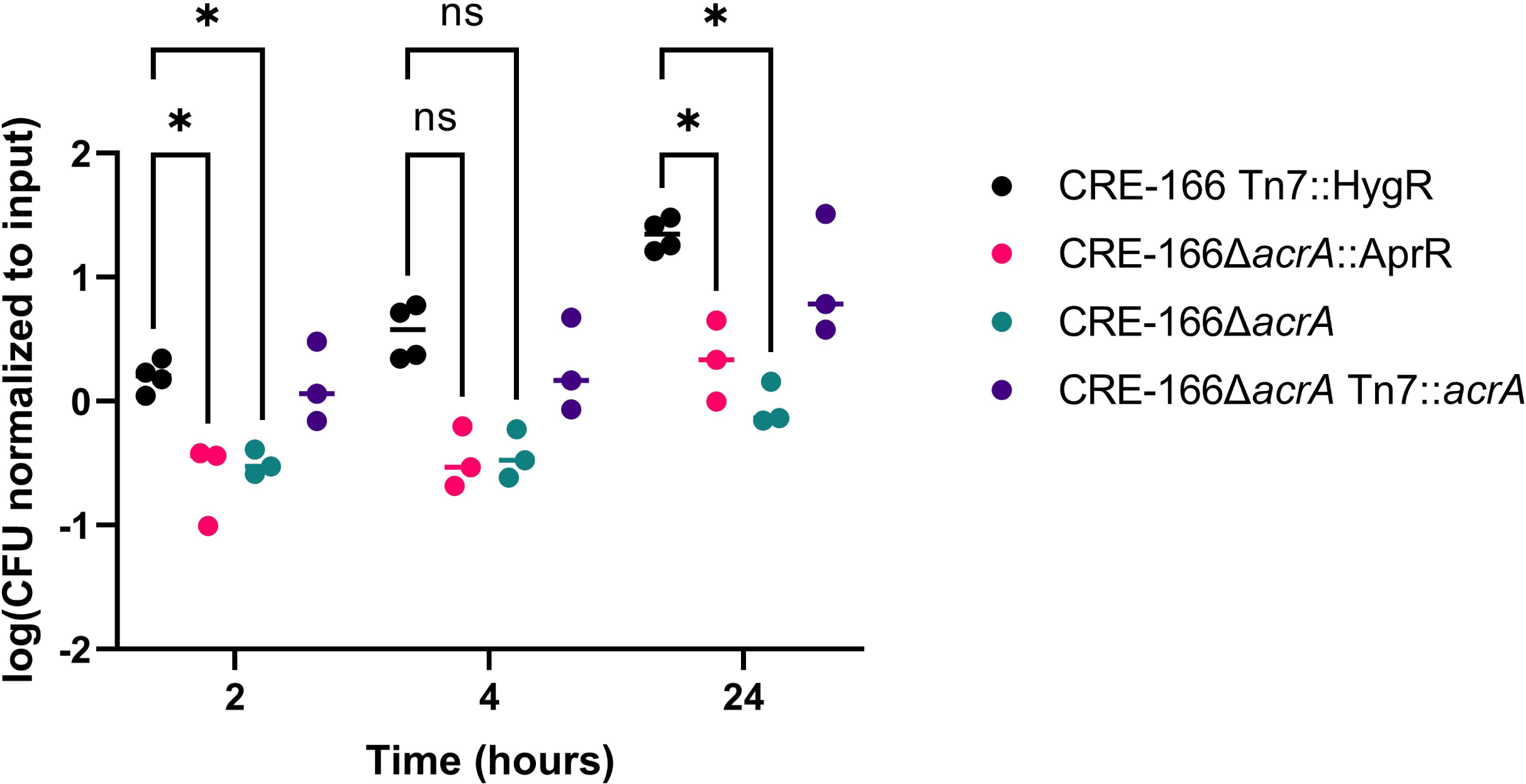
Resistance of CRE-166 and *acrA* mutants to 10% ox bile. Strains were inoculated into 10% ox bile (w/v), incubated, and CFU were plated for enumeration at the indicated timepoints. n = 3 biological replicates. Line denotes median. * indicates p < 0.05 in two-way ANOVA with Tukey’s HSD test.

## DISCUSSION

In this study, we aimed to answer the following question: does the genetic diversity of *K. pneumoniae* affect the colonization strategies of different strains? We compared three clinically-relevant strains (a globally distributed ST258 strain with a carbapenemase gene, an epidemic ST45 strain with an ESBL gene, and a non-epidemic antibiotic-susceptible strain) and identified a core set of genes used by all three strains to colonize the GI tract. However, we found other genes and pathways unique to one or two strains, highlighting the diversity of colonization strategies in this genetically diverse species. In particular, we confirmed the importance of three different factors (*acrA*, *carAB, tatABCD*) in colonization for all three strains to validate our screens.

Our results defined a core colonization program of 27 genes utilized by all 3 strains. Most of these genes (16 of 27) were related to metabolism, as expected since metabolic adaption to the anaerobic colon is a prerequisite for successful colonization. In addition, we also identified three genetic loci involved in pyrimidine and purine synthesis (*carAB* and *purC*/*purH*, respectively). Beyond metabolic genes, all strains relied on *yeiE*, which encodes a transcriptional activator. The genes regulated by YeiE in *K. pneumoniae* are not well characterized, but in *Salmonella enterica* this activator also controls GI colonization^19^, targeting flagellar genes (which *K. pneumoniae* does not possess). In addition, *tatA* and *tatC*, which encode components of the Tat folded-protein secretion apparatus, were identified as elements of the core colonization program, suggesting secreted factors may contribute to colonization, as they do for several gut pathogens^20–23^. In particular, the Tat-secreted peptidoglycan amidases, AmiA and AmiC, are necessary for colonization by *S. typhimurium*^24^, and *amiC* was identified in our CRE-166 screen (Supplemental Table 2). However, disruption of the Tat system also destabilizes the cell envelope^25^, decreasing resistance to bile acids^20^. Genes encoding the porin OmpC (*ompC*) and components of the Tol-Pal system (*tolA* and *pal*) were also identified as critical for colonization for all 3 strains. In addition to allowing diffusion of small solutes, OmpC is responsible for maintaining outer membrane leaflet asymmetry^26^. In a different fashion, the Tol-Pal system also aids in maintaining the integrity of the outer membrane^27^. Deletion of *pal* in *K. pneumoniae* increases sensitivity to bile, one of the host-derived stresses encountered in the GI tract^28^. A few other genes implicated in bile resistance (*cvpA* and *acrA*) were critical for colonization. In summary, the core GI colonization program of *K. pneumoniae* is composed of genes involved in energy generation, nucleotide biosynthesis, protein secretion, membrane homeostasis, and bile resistance.

We found many genes that contributed to colonization by one or two strains but not by all three strains. For instance, pathway analysis revealed that unlike the other two strains, Z4160 colonization factors were enriched for alanine, aspartate, and glutamate metabolism pathways. In addition, we found that CRE-166 depends on the sucrose porin *scrY* for colonization whereas Z4160 and KPN46 do not. It is possible that redundancy in either sucrose uptake or functional redundancy in other metabolic pathways can compensate for *scrY* deletion in Z4160 and KPN46 but not CRE-166. These findings support our hypothesis that colonization strategies differ between strains of *K. pneumoniae*.

The identification of *acrA* as necessary for GI colonization in all three strains has translational implications. AcrA is the periplasmic subunit of the tripartite efflux pumps that contain TolC and AcrB or AcrD^17^. In *E. coli* and *S. enterica*, these pumps export a large variety of substrates, including multiple classes of antibiotics and bile acids^17,29^. We demonstrated that the *acrA* mutant in CRE-166 was more susceptible to bile (Figure 7), suggesting a similar function in *K. pneumoniae*. Furthermore, the *acrA* deletion mutants were undetectable at 14 days post-inoculation in most of our competition experiments (Figure 6A-C). Because of its role in antibiotic resistance, the AcrAB efflux pump has been extensively studied, and several small molecule inhibitors are in varying stages of pre-clinical development^30^. We postulate that these inhibitors may have efficacy in preventing or eradicating *K. pneumoniae* GI colonization.

CRE-166 belongs to the globally disseminated ST258 group of carbapenemase-producing strains, and there is substantial interest in understanding how these strains successfully colonize patients. Jung and colleagues previously performed transposon insertion sequencing on a different ST258 strain, MH258, and our results have similarities. First, both studies found that disruption of genes involved in maltose metabolism conferred a fitness advantage in the GI tract. The lambda phage uses maltoporin LamB as its receptor^31^, so it is possible that the maltose system serves as a receptor for a lytic phage, bacteriocin, or other bacterial competition systems. Second, 9 of 35 MH258 colonization factors were also important for CRE-166. However, we also found 150 genes to be important for GI colonization with CRE-166 that were not identified for MH258. These differences may reflect genetic variation between the ST258 strains or differences in screen analysis and antibiotic regimens.

Our study had several limitations. All transposon insertion sequencing screens have a propensity to miss secreted factors because of trans-complementation. This may be one reason why we did not detect Type VI secretion systems in our screens even though they have been shown to contribute to colonization^32,33^. Interestingly, we did detect *amiC*, which encodes for a Tat-secreted substrate, but this substrate may remain localized within bacteria. Additionally, our screen was performed on Day 3 of colonization—the latest timepoint at which we did not observe a substantial bottleneck. Factors necessary for colonization on Day 3 may be different from those necessary for longer term colonization. However, these factors would likely be relevant for the development of prophylactic therapies against colonization, and our mutant studies show that deletion of several genes required for Day 3 colonization were also necessary at Day 14 (Figure 6). Additionally, we found that one strain-specific factor, *scrY*, had a stronger phenotype in targeted deletion and validation than another factor, *hha*. This may be explained by outlier read counts that have an inflated effect on the averages used to calculate log fold change. This type of technical limitation of the analysis may explain why *tolC,* which is usually complexed to *acrA,* was identified as a colonization factor for CRE-166 but not the two other strains. Another limitation is that we used as our input pool the mutant library that was inoculated into mice rather than a mutant library passaged in LB. As a result, some of the genes we identified as necessary for GI colonization were likely necessary for normal bacterial growth in rich medium. Such genes, although mechanistically less interesting, are in the strictest sense still necessary for colonization. Our study used only 3 strains, all of which were classical strains. The conserved genome between these clinical strains was 4558 CDS, which is larger than the core content between larger collections of *K. pneumoniae* strains (around 1700 CDS). Additional studies will be necessary to determine whether our findings can be extrapolated to a larger number of *K. pneumoniae* strains, including hypervirulent strains. Finally, our studies were done with C57BL/6 mice; experiments with different mouse strains are necessary to determine whether mouse background affects the genes necessary for colonization.

In conclusion, our study found that while different strains of *K. pneumoniae* rely on different genes and pathways to colonize the GI tract, there is a core set of colonization factors used by multiple strains. Inhibition of the proteins encoded by shared genes could theoretically block colonization by nearly all *K. pneumoniae* strains, whereas inhibition of lineage-specific factors could selectively block colonization by specific genotypes.

## MATERIALS AND METHODS

### Bacterial strains and cultures

CRE-166, KPN46, and Z4160 are *K. pneumoniae* clinical isolates from Northwestern Memorial Hospital in Chicago collected between 2014 and 2015. CRE-166 and KPN46 were previously described^34,35^, whereas Z4160 was first used in the current study. *E. coli* strain PIR1 was used for cloning, and *E. coli* β3914 (diaminopimelic acid auxotroph) was used to mate plasmids into *K. pneumoniae*^36,37^.

Bacteria were grown in LB 37°C unless otherwise stated. When appropriate, the following antibiotics were added: carbenicillin (100 μg/mL), hygromycin (100 μg/mL), or apramycin (50 μg/mL). Medium for β3914 was supplemented with 10 μg/mL of DAP.

### Preparation of complete genomes

Genomic DNA was extracted from CRE-166, KPN46, and Z4160 and sequenced on Illumina and Nanopore platforms to create complete genomes. Annotation was performed using the NCBI Prokaryotic Genome Annotation Pipeline (Z4160)^38^ or Prokka (CRE-166 and KPN46)^39^.

### Identification of shared genes

To identify genes shared between the three *K. pneumoniae* strains, we used the program Spine and defined shared coding sequences as those with >85% sequence homology between strains^40^. We defined the core genome of *K. pneumoniae* as sequences present in 95% of 323 strains from a previous study^4^ (with homologous genes defined as those with >85% similarity between strains).

### Murine model of GI Colonization

Six- to eight-week-old C57BL/6 mice (Jackson Laboratories) received 5 daily intraperitoneal injections of vancomycin (350 mg/kg, Hospira) unless otherwise indicated. For gavage with individual strains, inocula of 10^8^ CFU in 50 μl of PBS were used. Mice received daily cage changes to minimize coprophagy. CFU were enumerated by homogenization of fecal pellets in PBS with the Benchmark Bead Blaster 24 (Benchmark Scientific) followed by serial dilution and plating on LB agar with carbenicillin.

Transposon insertion sequencing experiments were performed as above. Frozen aliquots of the transposon libraries were revived for 2 hours in 25 mL of LB. CFU in fecal pellets were quantified as above, and DNA was extracted from the homogenates with the Maxwell 16 system.

For competitive colonization experiments, inocula of 1:1 mixtures of a hygromycin-resistant parental strain and an isogenic apramycin-resistant mutant (10^8^ CFU each) were created. Fecal CFU burdens were enumerated as above by plating on LB agar with hygromycin or apramycin. Competitive indices (CIs) were calculated as the ratio of mutant CFU to parent strain CFU, normalized to the input ratio.

Mice were housed in a containment ward of the Center for Comparative Medicine at Northwestern University. Experiments were approved by the Northwestern University Institutional Animal Care and Use Committee in compliance with ethical regulations.

### Construction of transposon mutant libraries

A suicide plasmid suitable for Himar1 mariner transposon mutagenesis in highly antibiotic-resistant strains was generated from pSAM*erm* (gift from G. Pier^41^). Briefly, pSAM*erm* was modified by replacement of the erythromycin-resistance cassette with a hygromycin-resistance cassette (HygR), and the MmeI sites in HygR were removed by site-directed mutagenesis. The resulting plasmid, pSAM*hygSDM*, was transformed into *E. coli* β3914 to generate a donor strain for conjugation with *K. pneumoniae* recipient strains. To select for transconjugants and eliminate β3914, we plated on LB agar supplemented with hygromycin (but lacking DAP) and incubated at 37°C overnight. Colonies were scraped and resuspended in LB with 25% glycerol. The resulting library was stored at −80°C.

### Arbitrary PCR for library quality control

Genomic DNA was extracted from 32 randomly selected colonies from each library, and two rounds of nested PCR were performed to amplify the transposon insertion site for subsequent Sanger sequencing. Primer sequences are listed in Supplemental Table 3.

### Preparation of DNA for transposon insertion sequencing

DNA extracted from fecal pellets was prepared for insertion site sequencing using the method of Kazi and colleagues^42^. Three replicates for each input (technical) and output (biological) pool were prepared, plus one technical replicate of an output sample. DNA was sheared to ∼250 bp fragments with the E220 ultrasonicator (Covaris), and insertion sites were amplified with addition of capture/sequencing sites and barcodes. Final library pool concentrations were quantified with Kapa library quantification (Roche) and sequenced on an Illumina MiSeq.

### Transposon insertion sequencing data analysis

A modified version of the previously described ESSENTIALS pipeline^11^ was used to identify genes necessary for growth in LB and genes that contributed to colonization^43^.

### Pathway analysis

KEGG identifiers were assigned to all genes using BlastKOALA, and KEGG pathways were assigned using KEGG Mapper^44,45^. A hypergeometric test was conducted in R (v4.2.2) to determine which KEGG pathways were enriched for colonization factors.

### Isogenic mutant construction

Deletion mutants were created using lambda red recombination as previously described^46^. All mutants were confirmed by whole-genome sequencing on Illumina platforms.

### Bile assays

Dehydrated ox bile was resuspended in water (10% w/v, Sigma) and filtered through a 0.2 μm filter. Strains were grown to an OD_600_ of 1.0, and 1 mL of each culture was pelleted and resuspended in 1 mL PBS. Each strain was inoculated (100 μl) into 900 μl of 10% ox bile, PBS, or LB and incubated with shaking at 37°C. At 0, 2, 4, and 24 hours, aliquots were removed and plated on LB agar for CFU enumeration.

### Data availability

The complete genomes of CRE-166, KPN46, and Z4160 have been deposited, and respective accession numbers are: GCA_016797255.2, GCA_021272285.2, GCA_030019695.1.

## ACKNOWLEDGEMENTS

We thank Mark Mandel for the β3914 strain and David Amici for comments on the manuscript. Support for this work was provided by the National Institutes of Health grants R01 AI118257, U19 AI135964, K24 AI04831, R21 AI164254, R21 AI153953 (A.R.H) and T32GM008152 (B.H.C), American Cancer Society grant 134251-CSDG-20-053-01-MPC (K.E.R.B), and American Heart Association grant 837089 (T.K.).

## BIBLIOGRAPHY

1. Giannella M, Trecarichi EM, De Rosa FG, et al. Risk factors for carbapenem-resistant *Klebsiella pneumoniae* bloodstream infection among rectal carriers: a prospective observational multicentre study. Clinical Microbiology and Infection 2014;20:1357–62.

2. Gorrie CL, Mirčeta M, Wick RR, et al. Gastrointestinal Carriage Is a Major Reservoir of *Klebsiella pneumoniae* Infection in Intensive Care Patients. Clinical Infectious Diseases: An Official Publication of the Infectious Diseases Society of America 2017;65:208.

3. Lam MMC, Wick RR, Watts SC, Cerdeira LT, Wyres KL, Holt KE. A genomic surveillance framework and genotyping tool for *Klebsiella pneumoniae* and its related species complex. Nature Communications 2021;12:1–16.

4. Holt KE, Wertheim H, Zadoks RN, et al. Genomic analysis of diversity, population structure, virulence, and antimicrobial resistance in *Klebsiella pneumoniae*, an urgent threat to public health. Proceedings of the National Academy of Sciences of the United States of America 2015;112:E3574.

5. Jung H-J, Littmann ER, Seok R, et al. Genome-Wide Screening for Enteric Colonization Factors in Carbapenem-Resistant ST258 *Klebsiella pneumoniae*. mBio 2019;10.

6. Maroncle N, Balestrino D, Rich C, Forestier C. Identification of *Klebsiella pneumoniae* genes involved in intestinal colonization and adhesion using signature-tagged mutagenesis. Infection and Immunity 2002;70:4729–34.

7. Struve C, Forestier C, Krogfelt KA. Application of a novel multi-screening signature-tagged mutagenesis assay for identification of *Klebsiella pneumoniae* genes essential in colonization and infection. Microbiology 2003;149:167–76.

8. Cerqueira GC, Earl AM, Ernst CM, et al. Multi-institute analysis of carbapenem resistance reveals remarkable diversity, unexplained mechanisms, and limited clonal outbreaks. Proceedings of the National Academy of Sciences of the United States of America 2017;114:1135.

9. Baggs J, Fridkin SK, Pollack LA, Srinivasan A, Jernigan JA. Estimating National Trends in Inpatient Antibiotic Use Among US Hospitals From 2006 to 2012. JAMA Internal Medicine 2016;176:1639–48.

10. Administration. UFD. Estimating the Maximum Safe Starting Dose in Initial Clinical Trials f. 2018.

11. Zomer A, Burghout P, Bootsma HJ, Hermans PWM, van Hijum SAFT. ESSENTIALS: Software for Rapid Analysis of High Throughput Transposon Insertion Sequencing Data. PLOS ONE 2012;7:e43012.

12. Jana B, Cain AK, Doerrler WT, et al. The secondary resistome of multidrug-resistant *Klebsiella pneumoniae*. Scientific Reports 2017;7.

13. Ramage B, Erolin R, Held K, et al. Comprehensive Arrayed Transposon Mutant Library of *Klebsiella pneumoniae* Outbreak Strain KPNIH1. J Bacteriol 2017;199.

14. Bachman MA, Breen P, Deornellas V, et al. Genome-Wide Identification of Klebsiella pneumoniae Fitness Genes during Lung Infection. MBio 2015;6:e00775.

15. Charlier D, Nguyen Le Minh P, Roovers M. Regulation of carbamoylphosphate synthesis in *Escherichia coli*: an amazing metabolite at the crossroad of arginine and pyrimidine biosynthesis. Amino Acids 2018;50:1647–61.

16. Palmer T, Berks BC. The twin-arginine translocation (Tat) protein export pathway. Nature Reviews Microbiology 2012;10:483–96.

17. Li X-Z, Plésiat P, Nikaido H. The Challenge of Efflux-Mediated Antibiotic Resistance in Gram-Negative Bacteria. Clinical Microbiology Reviews 2015;28:337.

18. Thanassi DG, Cheng LW, Nikaido H. Active efflux of bile salts by *Escherichia coli*. J Bacteriol 1997;179:2512.

19. Westerman TL, McClelland M, Elfenbein JR. YeiE Regulates Motility and Gut Colonization in *Salmonella enterica* Serotype *Typhimurium*. mBio 2021;12.

20. Reynolds MM, Bogomolnaya L, Guo J, et al. Abrogation of the twin arginine transport system in *Salmonella enterica* serovar *Typhimurium* leads to colonization defects during infection. PloS One 2011;6:e15800.

21. Zhang L, Zhu Z, Jing H, et al. Pleiotropic effects of the twin-arginine translocation system on biofilm formation, colonization, and virulence in *Vibrio cholerae*. BMC Microbiology 2009;9:114.

22. Lavander M, Ericsson SK, Bröms JE, Forsberg Å. The Twin Arginine Translocation System Is Essential for Virulence of *Yersinia pseudotuberculosis*. Infection and Immunity 2006;74:1768.

23. Rajashekara G, Drozd M, Gangaiah D, Jeon B, Liu Z, Zhang Q. Functional characterization of the twin-arginine translocation system in *Campylobacter jejuni*. Foodborne Pathogens and Disease 2009;6:935–45.

24. Fujimoto M, Goto R, Hirota R, et al. Tat-exported peptidoglycan amidase-dependent cell division contributes to *Salmonella Typhimurium* fitness in the inflamed gut. PLoS Pathogens 2018;14.

25. Stanley NR, Findlay K, Berks BC, Palmer T. *Escherichia coli* Strains Blocked in Tat-Dependent Protein Export Exhibit Pleiotropic Defects in the Cell Envelope. J Bacteriol 2001;183:139.

26. Chong Z-S, Woo W-F, Chng S-S. Osmoporin OmpC forms a complex with MlaA to maintain outer membrane lipid asymmetry in *Escherichia coli*. Molecular Microbiology 2015;98:1133–46.

27. Szczepaniak J, Press C, Kleanthous C. The multifarious roles of Tol-Pal in Gram-negative bacteria. FEMS Microbiology Reviews 2020;44:490.

28. Hsieh P-F, Liu J-Y, Pan Y-J, et al. *Klebsiella pneumoniae* Peptidoglycan-Associated Lipoprotein and Murein Lipoprotein Contribute to Serum Resistance, Antiphagocytosis, and Proinflammatory Cytokine Stimulation. Journal of Infectious Diseases 2013;208:1580–9.

29. Nishino K, Latifi T, Groisman EA. Virulence and drug resistance roles of multidrug efflux systems of *Salmonella enterica* serovar *Typhimurium*. Molecular Microbiology 2006;59:126–41.

30. Sharma A, Gupta VK, Pathania R. Efflux pump inhibitors for bacterial pathogens: From bench to bedside. Indian Journal of Medical Research 2019;149:129.

31. Randall-Hazelbauer L, Schwartz M. Isolation of the Bacteriophage Lambda Receptor from *Escherichia coli*. J Bacteriol 1973.

32. Calderon-Gonzalez R, Lee A, Lopez-Campos G, et al. Modelling the Gastrointestinal Carriage of *Klebsiella pneumoniae* Infections. MBio 2023;e0312122.

33. Merciecca T, Bornes S, Nakusi L, et al. Role of *Klebsiella pneumoniae* Type VI secretion system (T6SS) in long-term gastrointestinal colonization. Scientific Reports 2022;12:16968.

34. Bulman ZP, Krapp F, Pincus NB, et al. Genomic Features Associated with the Degree of Phenotypic Resistance to Carbapenems in Carbapenem-Resistant *Klebsiella pneumoniae*. mSystems 2021;6.

35. Kochan TJ, Nozick SH, Medernach RL, et al. Genomic surveillance for multidrug-resistant or hypervirulent *Klebsiella pneumoniae* among United States bloodstream isolates. BMC Infectious Diseases 2022;22.

36. Wang N, Ozer EA, Mandel MJ, Hauser AR. Genome-Wide Identification of *Acinetobacter baumannii* Genes Necessary for Persistence in the Lung. mBio 2014;5.

37. Le Roux F, Binesse J, Saulnier D, Mazel D. Construction of a *Vibrio splendidus* Mutant Lacking the Metalloprotease Gene *vsm* by Use of a Novel Counterselectable Suicide Vector. Applied and Environmental Microbiology 2007;73:777.

38. Tatusova T, DiCuccio M, Badretdin A, et al. NCBI prokaryotic genome annotation pipeline. Nucleic Acids Research 2016;44:6614.

39. Seemann T. Prokka: rapid prokaryotic genome annotation. Bioinformatics 2014;30:2068–9.

40. Ozer EA, Allen JP, Hauser AR. Characterization of the core and accessory genomes of *Pseudomonas aeruginosa* using bioinformatic tools Spine and AGEnt. BMC Genomics 2014;15:737.

41. Skurnik D, Roux D, Aschard H, et al. A comprehensive analysis of *in vitro* and *in vivo* genetic fitness of *Pseudomonas aeruginosa* using high-throughput sequencing of transposon libraries. PLoS Pathogens 2013;9:e1003582.

42. Kazi MI, Schargel RD, Boll JM. Generating Transposon Insertion Libraries in Gram-Negative Bacteria for High-Throughput Sequencing. JoVE (Journal of Visualized Experiments) 2020:e61612.

43. Ozer EA. egonozer/essentials_local: Version 2.1. 2.1 ed: Zenodo; 2023.

44. Kanehisa M, Sato Y, Morishima K. BlastKOALA and GhostKOALA: KEGG Tools for Functional Characterization of Genome and Metagenome Sequences. Journal of Molecular Biology 2016;428:726–31.

45. Kanehisa M, Sato Y, Kawashima M. KEGG mapping tools for uncovering hidden features in biological data. Protein Science : a Publication of the Protein Society 2022;31:47–53.

46. Huang T-W, Lam I, Chang H-Y, Tsai S-F, Palsson BO, Charusanti P. Capsule deletion via a λ-Red knockout system perturbs biofilm formation and fimbriae expression in *Klebsiella pneumoniae* MGH 78578. BMC Research Notes 2014;7:13.

